# A novel humanized mouse model to study mucosal HIV-1 transmission and prevention

**DOI:** 10.1101/2020.08.31.274167

**Authors:** Kanika Vanshylla, Kathrin Held, Tabea M. Eser, Henning Gruell, Kanika Jain, Daniela Weiland, Jan Muench, Berthold Gruettner, Christof Geldmacher, Florian Klein

## Abstract

Humanized mice have been critical for HIV-1 research, but are inefficient at mucosal HIV-1 transmission. We present a fetal tissue-independent model called CD34T+ with enhanced human leukocyte levels in the blood and improved T cell homing to the gut-associated lymphoid tissue. CD34T+ mice are highly permissive to intra-rectal HIV-1 infection, which can be prevented by infusion of broadly neutralizing antibodies. Therefore, CD34T+ mice provide a novel platform for mucosal HIV-1 transmission and prevention studies.

## Main

Despite immense progress in the treatment of HIV-1 infection, transmission of the virus continues due to the lack of a vaccine that elicits full protection^1^. Anti-retroviral pre-exposure prophylaxis (PrEP) has shown efficacy in preventing transmission^2^. However, strict daily dosage requirements and disappointing results from clinical trials in African women warrant the need to develop alternatives^3^. The pre-clinical *in vivo* evaluation of molecules for HIV-1 prevention has been mostly limited to non-human primates (NHPs), along with some studies in Bone marrow Liver Thymus (BLT) humanized mice^4^. NHPs are susceptible to infection via sexual/mucosal routes, but the system entails using recombinant Simian-HIV to infect primate immune cells^4^. In contrast, humanized mice harbor human immune cells, can be infected with lab-adapted or primary isolates of HIV-1 and have been useful for analysis of antiviral efficacy of antiretroviral drugs and monoclonal antibodies^5^. BLT mice with high human cell reconstitution in the gut-associated lymphoid tissue (GALT), have been the primary model for mucosal HIV-1 transmission studies^6^. However, making BLT mice requires cumbersome surgical implantation of human fetal tissues, the use of which is accompanied by both ethical and legal restrictions. This has resulted in an urgent need to develop new mouse models for studying HIV-1 prevention that do not rely on fetal tissue^7^.

A prominent fetal tissue-independent method to humanize mice is to inject cord blood-derived CD34+ human stem cells^5^. Besides one report of successful HIV-1 infection via the mucosa, most CD34 humanized mice have low permissiveness to mucosal HIV-1 infection, which has been attributed to low human cell reconstitution in the GALT^8,9^. Here, we present a new model for HIV-1 prevention, called CD34T+, that does not rely on fetal tissue but allows robust mucosal HIV-1 transmission. Tested in parallel to the established CD34 model, CD34T+ mice achieve high levels of human cell reconstitution in both blood and the GALT. This facilitates high rates of mucosal HIV-1 transmission and allows the use of CD34T+ mice as a small animal model for testing anti-HIV-1 molecules for HIV-1 prevention.

To generate CD34T+ mice, individual umbilical cord blood samples were processed and stored in two fractions: CD34+ hematopoietic stem cells (HSCs) and mononuclear umbilical cord blood cells (UCBCs) (Fig. 1a). For humanizing mice, newborn NOD-*Rag1*^*null*^ *IL2rg*^*null*^ (NRG) mice were irradiated and injected with CD34+ HSCs 4-6h later (Fig. 1a and Methods). At 12 weeks of age, peripheral blood from the mice was analyzed to measure human leukocytes and mice harboring more than 1 human CD4+ T cell/μl blood were considered humanized and termed CD34 mice (data not shown). To augment human cell reconstitution, CD34 mice (range of 13-40 weeks old) were injected via the intra-peritoneum with donor-matched UCBCs. The use of donor-matched UCBCs was done to reduce the incidence of Graft-versus-Host (GvH) response^10^. In addition, to promote homing of human T cells to the GALT, mice were administered four doses of 250 ng human interleukin-7 (IL-7)^11^ (Fig. 1a). The UCBC and IL-7 treated mice were termed CD34T+.

**Figure 1:**
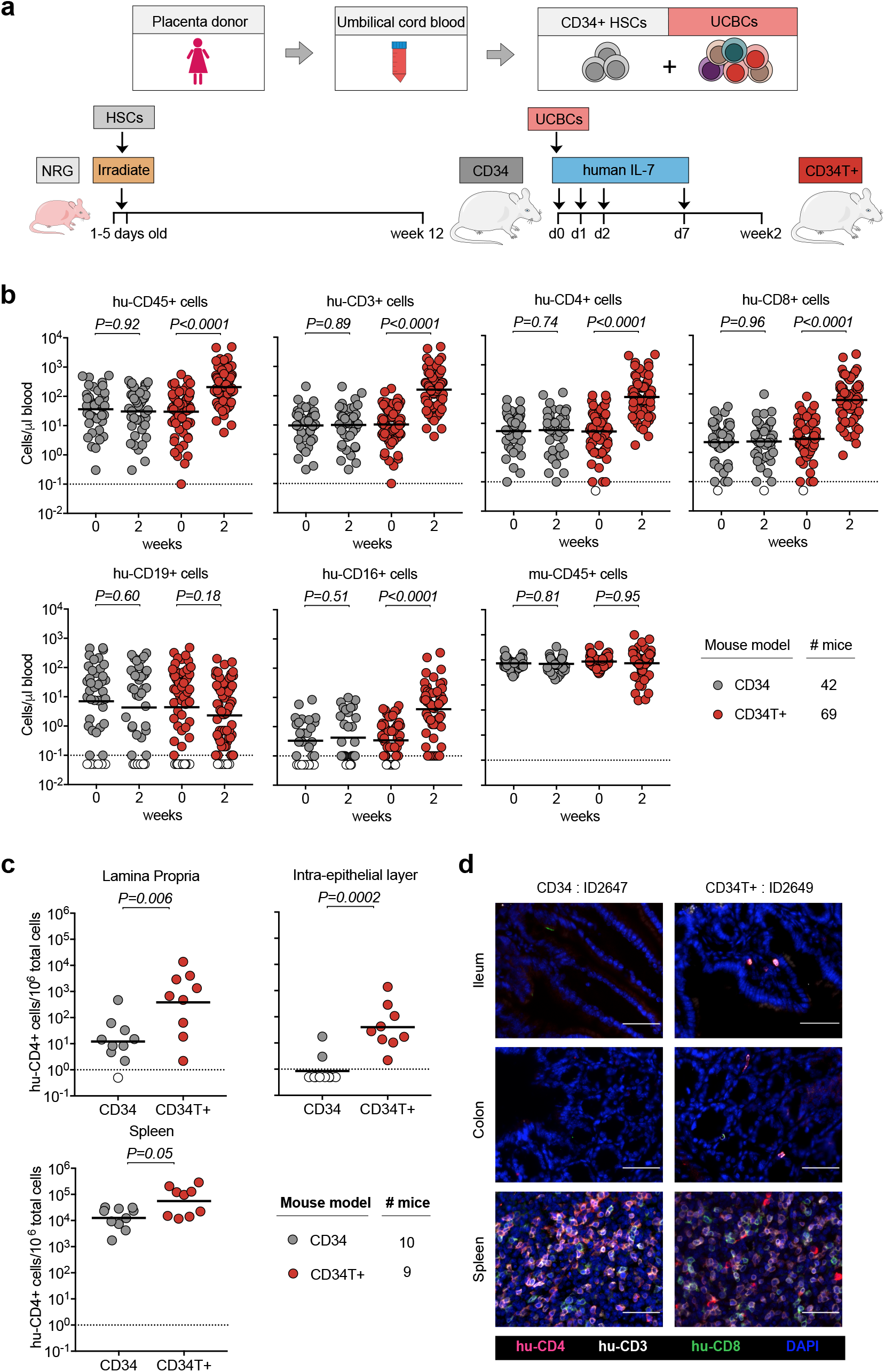
Generation of CD34T+ mice and reconstitution levels of human cells. **a**, Human umbilical cord blood from each donor was processed and stored as CD34+ HSCs and UCBCs. 1-5 days old NRG mice were subjected to sub-lethal irradiation and intra-hepatic injection of CD34+ HSCs. At 12 weeks of age, mice that had positively engrafted human leukocytes were termed CD34 mice. To obtain CD34T+ mice, CD34 mice were injected with donor-matched UCBCs and human-IL-7 at the indicated time-points. 2 weeks post UCBC injection, resulting mice were termed CD34T+. **b**, FACS analysis of leukocyte populations in the blood of CD34T+ mice before UCBC injection (week 0) and 2 weeks after UCBC injection (week 2) along with untreated CD34 mice analyzed in parallel. Points shown in white were below limit of detection of assay. Statistical analysis done using Wilcoxon matched-pairs signed rank test **c**, FACS analysis of CD34 and CD34T+ mice for the presence of human CD4+ T cells in the gut LP, IEL and the spleen at week 2. Points shown in white were below limit of detection of assay. Statistical analysis was done using Mann-Whitney U test. **d**, Immunohistochemical analysis of human T cells in the ileum, colon and spleen of CD34 and CD34T+ mice at week 2. Human CD3+ (white), CD4+ (red), CD8+ (green) cells were labeled for detection along with DAPI for visualizing the nucleus. Scale bar is 50 μm. 3 mice per group were analyzed with at least 3 different slices from different areas of each mouse tissue being analyzed. Representative images were selected for display.

We compared CD34T+ mice before (week 0) and after (week 2) UCBC injection, along with a control group of CD34 mice for changes in human leukocyte levels (FACS gating strategy in supplementary Fig. 1a). CD34T+ mice showed increased absolute levels of human CD45+ cells in the blood. The primary lymphocyte population to undergo expansion was CD3+ T cells and included both CD4+ and CD8+ T cell populations. In addition, we observed a significant increase in CD16+ cells in CD34T+ mice, while CD19+ B cells and mouse CD45+ leukocytes did not show significant change (Fig. 1b, supplementary Fig. 1b and Table S1). In the CD34 control group, levels of all analyzed human and mouse cell populations remained similar over time (Fig. 1b, supplementary Fig. 1b and Table S2). Our analysis encompassed 18 umbilical cord blood donors, confirming that CD34T+ mice can be successfully generated from different donors (supplementary Fig. 1c). These data show that the CD34T+ mice exhibit high T cell reconstitution in the peripheral blood and achieve levels which are comparable to cell populations in human blood^12^.

Next, we examined human cell reconstitution in lymphoid tissues by purifying cells from spleen, gut lamina propria (LP) and gut intra-epithelial layer (IEL) (supplementary Fig. 2a). CD34T+ mice exhibited significantly higher levels of human CD4+ T cells in the LP and IEL than CD34 mice (Fig. 1c). CD34T+ mice also had more CD4+ T cells in the spleen but this difference was not statistically significant (Fig. 1c). Similar levels of murine CD45+ cells confirmed similar purification efficiency across mice from both groups (supplementary Fig. 2b). To verify that analyzed cells did not originate from tissue-draining blood vessels, we performed immunohistochemical (IHC) staining of T cell distribution in intact tissue. IHC also confirmed the presence of human T cells in gut sections from CD34T+ mice, but none in the sections from CD34 mice (Fig. 1d), where FACS analysis of whole gut tissue had also only revealed a scarce human CD4+ T cell population. Human CD4+ T cell levels in the blood vs. lymphoid tissues revealed a positive linear relationship with Pearson *r* values of 0.705, 0.782 and 0.827 for LP, IEL and spleen respectively (supplementary Fig. 2c). Thus, the high human CD4+ T cell levels in blood of CD34T+ mice are accompanied by efficient homing of human T cells to the GALT.

We expected low mucosal HIV-1 transmission in CD34 mice, based on the low human T cell reconstitution in the GALT (Fig. 1). As expected, intra-rectal (i.r.) challenge of CD34 mice (supplementary Fig. 3a) with NL4-3_YU2_ HIV-1 resulted in viremia in only 4 out of the 15 challenged mice (supplementary Fig. 3b). Of note, all mice were confirmed to be susceptible to HIV-1 since subsequent intra-peritoneal (i.p.) challenge of the remaining 11 mice resulted in viremia (supplementary Fig. 3c). To identify the tissue reservoir of infected human CD4+ T cells after i.p. infection in CD34 mice, we used RNAscope to visualize HIV-1 RNA in spleen and GALT (ileum and colon) sections. We found infected cells in the spleen, but none in the GALT of the CD34 mice (supplementary Fig. 3d).

In contrast to CD34 mice, in CD34T+ mice, i.r. challenge with the same NL4-3_YU2_ HIV-1 stock all mice became infected, exhibiting 100% success rate (Fig. 2a, 2b and 2c). Moreover, mean plasma viremia of ∽10^6^ HIV-1 RNA copies/ml at week 3 in CD34T+ mice (Fig. 2d) is comparable to early stages of infection in humans^13^. HIV-1 RNAscope staining revealed infected human CD4+T cell reservoirs in both GALT and spleen sections from CD34T+ mice (Fig. 2e). Single genome sequencing (SGS) and phylogenetic analysis of plasma virions revealed similar degree of *in vivo* viral diversification following i.r. challenge in CD34T+ mice and i.p. challenge in CD34 mice (supplementary Fig. 4a-c). Thus, CD34T+ is a highly effective humanized mouse model to study mucosal HIV-1 transmission.

**Figure 2:**
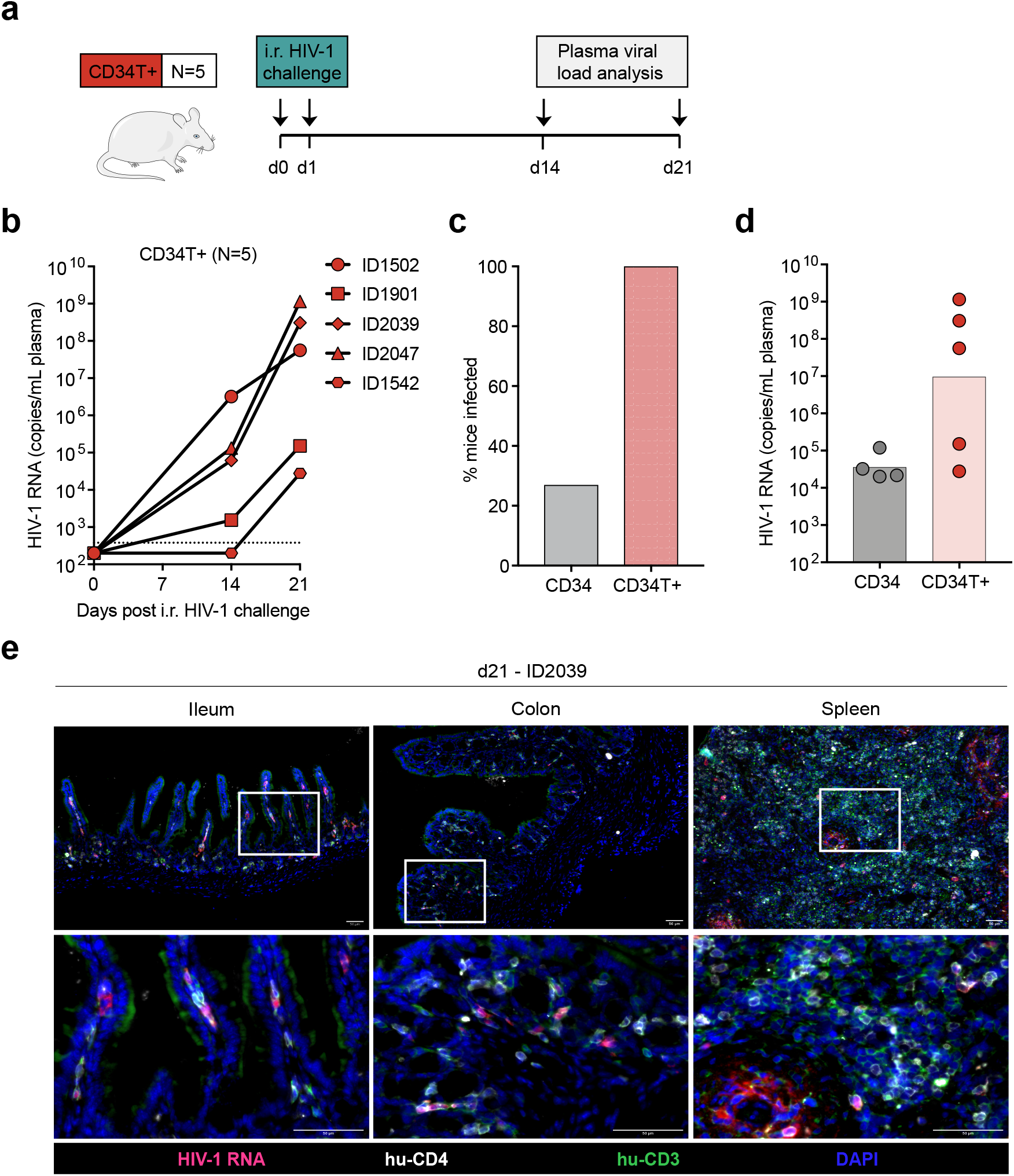
Mucosal transmission of HIV-1 in CD34T+ mice. **a**, Schematic representation of the experiment design. CD34T+ control mice were given i.r. challenges with NL4-3YU2 HIV-1 on consecutive days and bled at the indicated time points for viral load measurements. **b**, Plasma HIV-1 RNA levels in the blood of CD34T+ mice after intra-rectal challenge with NL4-3_YU2_ HIV-1. The dotted line represents the quantification limit of 384 HIV-1 RNA copies/ml of the assay. **c**, Comparison of infectivity rates (in %) of CD34 and CD34T+ mice post i.r. HIV-1 challenge. **d**, Plasma HIV-1 RNA levels at week 3 after i.r. challenge with HIV-1 of CD34 and CD34T+ mice. **e**, RNA-scope detection of HIV-1 infected cells in the ileum, colon and spleen of CD34T+ mice harvested at day 21 post i.r. HIV-1 challenge. HIV-1 RNA (pink), CD4+ (white), CD3+ (green) cells were labeled for detection along with DAPI for visualizing the nucleus. Scale bar is 50 μm. 3 mice per group were analyzed with 3 different slices from each mouse tissue being analyzed. Representative images were selected for display. i.r. intra-rectal

One main goal behind developing CD34T+ mice was to have a mucosal HIV-1 prevention model to rapidly evaluate novel anti-HIV-1 molecules, for instance, broadly neutralizing antibodies (bNAbs). Antibody-mediated prevention offers advantages like long systemic half-life, minimal side-effects and effector functions that include activation of other immune cells^14^. To test prevention in CD34T+ mice, we used a tri-mix combination of three potent bNAbs, 3BNC117, 10-1074 and SF12^15-17^. These antibodies target different epitopes on the HIV-1 envelope and are potent against NL4-3_YU2_ HIV-1 (supplementary Fig. 5a and 5b). The control group received i.r. HIV-1 challenges, whereas the tri-mix group received an i.p. infusion of the antibodies 1 day before the first i.r. HIV-1 challenge (Fig. 3a). As before, all control group mice became infected within 3 weeks post i.r. HIV-1 challenge (Fig. 3b). In contrast, all tri-mix group mice were protected from HIV-1 infection and remained aviremic up to 8 weeks post HIV-1 challenge despite plasma antibody levels dropping to undetectable levels (Fig. 3b, 3c and supplementary Fig. 5c, 5d and 5e). This proves that CD34T+ mice can be used for evaluating the protective function of anti-HIV-1 antibodies.

**Figure 3:**
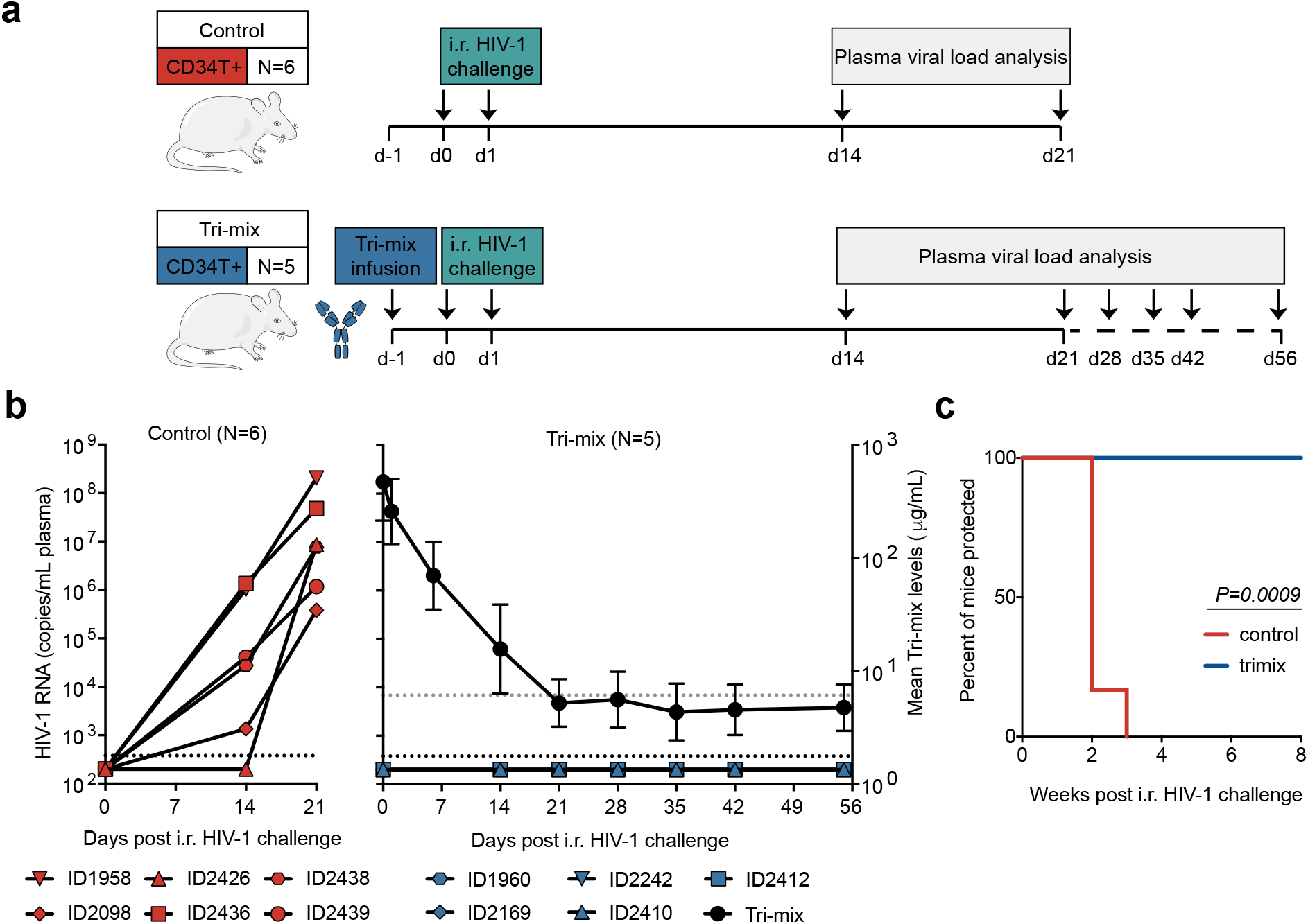
Antibody-mediated prevention of mucosal HIV-1 infection in CD34T+ mice. **a**, Schematic representation of the experimental design. CD34T+ control mice were given high dose i.r. challenges with NL4-3_YU2_ HIV-1 on consecutive days, whereas, the CD34T+ tri-mix mice additionally received i.p. injection of 2 mg each of 3BNC117, 10-1074 and SF12 one day before the 1^st^ HIV-1 challenge. **b**, Plasma HIV-1 RNA levels over time in the control and tri-mix mice. The lower black dotted line represents the assay quantification limit of 384 HIV-1 RNA copies/ml plasma. The right Y-axis for the tri-mix mice shows mean (N=5) neutralizing antibody tri-mix levels in plasma over time. The upper grey dotted line represents the limit of detection of the antibody measurement assay. **c**, Kaplan-Meier curve depicting rate of protection from mucosal HIV-1 infection in control and tri-mix mice from (**b**) statistical analysis done with Log-rank (Mantel-Cox) test. i.r. intra-rectal

In conclusion, CD34T+ mice demonstrate high peripheral and tissue reconstitution with human cells and the homing of T cells to the GALT. High T cell reconstitution in the GALT makes an ideal tissue environment for mucosal HIV-1 transmission in the CD34T+ mice. One limitation of CD34T+ mice is the onset of anemia in some animals. Development of anemia due to human phagocyte mediated erythrocyte depletion has been reported in humanized MISTRG mice^18^. It is possible that depletion of phagocytes in CD34T+ mice would help prevent anemia, without affecting the high transmission rates of HIV-1. Despite this, CD34T+ mice are a significant step forward towards our goal of having small animal models where complex processes like mucosal HIV-1 transmission can be recapitulated and molecules can be evaluated for their effectiveness in preventing HIV-1 infection.

Over the last years, bNAbs have emerged as a highly viable option for HIV-1 therapy and prevention^14^. However, due to the high cost burden of clinical trials, VRC01 is the only molecule that has entered human clinical trials for antibody-mediated prevention (NCT02716675 and NCT02568215). It is very likely that a globally effective prevention strategy will require a combination of highly potent antibodies^14^. Rapid *in vivo* testing of such combinations can further our goal of reducing the global HIV-1 burden. NHPs have been the dominant choice for such testing, but these studies incur high costs, resources and time, tipping the scale towards the use of small animal models. Although, HIV-1 prevention studies in BLT mice have been useful, the need for surgical expertise and access to highly restricted human fetal tissue complicates their large-scale use. The need for alternative models is highlighted by the recent notice from the National Institute of Health (NOT-OD-19-042) calling for development of fetal tissue-independent humanized mice. CD34T+ mice can be generated without complex surgery, are fetal tissue-independent and still achieve high human cell reconstitution. These attributes, along with the high efficiency of mucosal HIV-1 transmission, make CD34T+ mice a useful pre-clinical platform for testing of newly designed antibodies/drugs for HIV-1 prevention.

## Acknowledgements

We thank members of the Klein lab for helpful discussions; Hannah Jannicki, Julia Schmatz, Lukas Scheuengraber and Carola Ruping for technical assistance; staff of the Animal facility at Weyertal, University of Cologne to help maintaining the Humanized Mouse Core Cologne (HMCC); and the Cologne Center for Genomics (CCG) for support with sequencing. Imaging analysis of mouse tissue was partly funded by a grant of the Verein zur Förderung von Wissenschaft und Forschung an der Medizinischen Fakultät der Ludwig-Maximilians-Universität München e.V. (to K.H.). This work was mainly funded by grants from the German Center for Infection Research (DZIF; to F.K.), the German Research Foundation (CRC 1279 and CRC 1310 to F.K.), and the European Research Council (ERC-StG639961 to F.K.).

## Author contributions

Study was designed by F.K. and K.V.; Mouse experiments were performed by K.V. and H.G.; FACS analysis and viral load measurements were done by K.V.; Mouse tissue imaging was performed by K.H. and T.M.E.; Cord-blood samples were processed by K.J.; Help with mouse work was provided by D.W.; Resources were provided by H.G., B.G and J.M.; Supervision was provided by C.G. and F.K.; Manuscript was written by K.V. and F.K.; All authors have read the manuscript.

---

**Supplementary Figure 1: Analysis of leukocyte populations in the blood of CD34 and CD34T+ mice**

**a**, Gating strategy depicted with representative FACS plots from analysis of blood from a CD34T+ mouse ID1502 before (week 0) and after (week 2) UCBC and human IL-7 treatment. **b**, Mean cell numbers per μl blood and fold change between week 0 (w0) and week 2 (w2) from CD34 and CD34T+ mice shown in **Fig. 1b. c**, Analysis of human leukocyte expansion based on mean cell numbers per μl blood in CD34T+ mice (N=69) divided according to the cord blood donor.

**Supplementary Figure 2: Analysis of leukocyte populations in the tissue of CD34 and CD34T+ mice**

FACS analysis of CD34 and CD34T+ mice shown in Fig. 2c for the presence of human CD3+ T cell, human CD8+ T cells, human CD19+ B cells (**a**) and mouse CD45+ cells (**b**) in the gut LP, IEL and the spleen at week 2. Points shown in white were below limit of detection of assay. Statistical analysis done using Mann-Whitney U test. **c**, Combined Pearson correlation curves plotted for CD34 (N=10) and CD34T+(N=9) mice to analyze the relationship between human CD4+T cell levels in the blood vs. lymphoid tissue: LP, IEL and spleen).

**Supplementary Figure 3: Analysis of HIV-1 infection in CD34 mice**

**a**, Schematic representation of the experiment design. CD34 mice were given i.r. challenges with NL4-3_YU2_ HIV-1 on consecutive days and bled at the indicated time points for testing viremia. Mice that remained un-infected after i.r. challenge were challenged via i.p. injection of NL4-3_YU2_ HIV-1 to confirm susceptibility to infection. **b**, Plasma HIV-1 RNA levels in the blood of CD34 mice after i.r. challenge with high dose NL4-3_YU2_ HIV-1. **c**, Plasma HIV-1 RNA levels in CD34 mice after i.p. challenge with NL4-3_YU2_ HIV-1. The dotted line represents the detection limit of 384 HIV-1 RNA copies/ml of the assay in (**b**) and (**c**). **d**, RNA-scope detection of HIV-1 infection in the ileum, colon and spleen of CD34+ mice harvested at day 21 post i.p. HIV-1 challenge. HIV-1 RNA (pink), CD4+ (white), CD3+ (green) cells were labeled for detection along with DAPI for visualizing the nucleus. Scale bar is 50 μm. 3 mice per group were analyzed with 3 different slices from each mouse tissue being analyzed. Representative images were selected for display. i.r. intra-rectal, i.p. intra-peritoneal

**Supplementary Figure 4: Analysis of SGS-derived *env* sequences from plasma of HIV-1 infected mice**

**a**, Phylogenetic tree of SGS-derived *env* sequences from the plasma of CD34T+ mice and CD34 mice at day 21 post post i.r. or i.p. challenge with NL4-3_YU2_ HIV-1 respectively. **b**, Analysis of mutation rates at the amino acid and nucleotide level in the *env* gene of mice from **a. c**, Amino acid alignment of plasma SGS-derived *env* sequences from mice analyzed in (**a** and **b**). Red bars indicate amino acid residues with mutations relative to the wild-type NL4-3_YU2_ HIV-1 *env*. Amino acid numbering is based on HIV-1_YU2_ *env*. i.r. intra-rectal, i.p. intra-peritoneal

**Supplementary Figure 5: Neutralization potency of bNAb tri-mix against NL4-3**_**YU2**_ **HIV-1**

**a**, Neutralization potency of individual bNAbs (3BNC117, 10-1074 and SF12) or an equimolar tri-mix (combi.) against the NL4-3_YU2_ HIV-1 infectious molecular clone represented as 50% inhibitory concentration (IC50) or 80% inhibitory concentration (IC80) along with the maximum percent inhibition (MPI) values. **b**, Neutralization curve of the individual bNAbs along with the tri-mix measured in (**a**). **c**, Antibody concentration of the tri-mix in the plasma of CD34T+ tri-mix mice from Fig. 3 measured using the TZM-bl assay. **d**, 50% Inhibitory dilution (ID50) values of the tri-mix in the plasma of CD34T+ tri-mix mice from **Fig. 3** measured against the NL4-3_YU2_ HIV-1 infectious molecular clone. ID50 values at the time of i.r. HIV-1 challenge (d0 and d1) are highlighted in red. **e**, Tri-mix concentration in the plasma of CD34T+ tri-mix mice from **Fig. 3** measured against the NL4-3_YU2_ HIV-1 infectious molecular clone. Values at the time of i.r. HIV-1 challenge (d0 and d1) are highlighted in red and values below the detection limit of the assay are highlighted in grey.

## Methods

### Collection of umbilical cord blood and placenta derived cells

Placental and umbilical cord tissue were collected from donors who gave their written consent under a protocol approved by the Institutional Review Board of the University of Cologne (16-110). CD34+ HSCs were isolated from the cord blood and perfused placental tissue by using the CD34 Microbead Kit (Miltenyi Biotec) and stored at −150°C. The mononuclear cell fraction (UCBCs) of the cord blood was collected as part of the flow-through of the CD34 Microbead kit and stored at −150°C.

### Generation of humanized mice

NOD-*Rag1*^*null*^ *IL2rg*^*null*^ (NRG) mice were purchased from The Jackson Laboratory and bred and maintained at the Decentralized Animal Facility of the University of Cologne. Mice were housed under specific-pathogen-free conditions with 12-hour day/night conditions and given a diet of ssniff food. CD34 humanized mice were generated by engraftment with CD34+ HSCs isolated from umbilical cord blood as follows: 1-5 days old NRG mice were sub-lethally irradiated at a dose of ∽2.0 Gy and 4-6 hours later injected intra-hepatically with 2×10^5^ purified CD34+ HSCs. 12 weeks post engraftment, mouse blood was analyzed by FACS for the presence of human PBMCs. Mice that positively engrafted human CD4+ T cells were included in experiments and termed CD34 mice. For generating CD34T+ mice, on day 0, CD34 mice were injected subcutaneously with 250 ng human IL-7 (PeproTech) in PBS (Gibco), three hours before injection with donor-matched 30-45×10^6^ UCBCs derived from the same cord blood donor used for CD34+ HSC engraftment. Mice were administered 250 ng human IL-7 in PBS on day 1,2 and 7 post UCBC injection. At 2 weeks post UCBC injection, expansion of human leukocytes was analyzed by FACS and these mice were henceforth termed CD34T+ and used for further analysis/experiments. All mouse experiments were authorized by the State Agency for Nature, Environmental Protection and Consumer Protection North Rhine-Westphalia (LANUV).

### Isolation of cells from mouse spleen or gut-associated lymphoid tissue

Mice were sacrificed using cervical dislocation and the spleen, large intestine and small intestine were excised. All isolated cells or tissue were stored at 4°C in medium composed of RPMI 1640 medium GlutaMAX™ supplement (Thermo Fisher) containing 10% fetal bovine serum (FBS) (Sigma Aldrich) and 1% Penicillin/Streptomycin (Gibco) until further processing. The spleen was processed by homogenization of the tissue using a syringe piston on a 70 μm cell strainer (Corning) and passing the cell suspension through a 18G syringe needle. Erythrocytes were lysed with ACK lysis buffer (Thermo Fisher) and the splenic cells were washed in cell medium and analyzed by FACS. For isolation of lymphocytes from gut-associated lymphoid tissue, the intestine tissue was flushed with PBS. 1 cm dissected sections were pre-digested to break down the extracellular matrix by incubating twice at 37°C for 15 min in medium containing 25 mM EDTA followed by vortexing and passing the solution through a 70 μm cell strainer to obtain the intra-epithelial lymphocyte fraction. The rest of the tissue was digested by incubating twice at 37°C for 20 min in medium containing 1 mg/ml Collagenase D (Sigma-Aldrich), 0.1 mg/ml DnaseI (Sigma-Aldrich) and 0.5 mg/ml Dispase (Sigma-Aldrich), followed by vortexing and passing the solution through a 70 μm cell strainer (Corning) to obtain the lamina propria (LP) fraction. The IEL and LP fractions were washed in medium and passed through 70 μm and 40 μm cell strainers consecutively. The lymphocytes were purified by performing a density gradient using 30% (vol/vol) Histopaque solution (Sigma-Aldrich) and purified cells analyzed by FACS.

### Flow cytometry analysis of cell populations in mouse blood or tissue

Mice were bled from the submandibular vein and the erythrocytes in the blood were lysed with ACK lysis buffer and stained with the following antibodies: mouse-CD45-PECy7 (BioLegend; clone 30-F11), human-CD45-Pacific Orange (Thermo Fisher; clone HI30), human-CD19-APC (BD Pharmigen; clone HIB19), human-CD3-Pacific Blue (BD Pharmigen; clone UCHT1), human-CD4-PE (BD Pharmigen; clone L120), human-CD8-FITC (BD Pharmigen; clone SK1), human-CD16-AF700 (BD Pharmigen; clone CD2F1). Samples were washed and analyzed in FACS Buffer (PBS containing 2% FBS and 2 mM EDTA pH=8.0). Mouse tissue lymphocytes were isolated as detailed above and 1×10^6^ cells were used for FACS staining. FACS data were acquired using a BD FACSAria Fusion flow cytometer and sample analysis was done on the FlowJo v10.2 software with final graphs and statistical analysis being performed in GraphPad Prism 7.

### Production of replication competent HIV-1 using 293-T cells

Replication-competent NL4-3_YU2_^19^ HIV-1 molecular clone was used to transfect HEK 293T cells using the FuGENE 6 Transfection Reagent (Promega). The virus culture supernatant was harvested at 48 hours post transfection and stored at −80°C or −150°C.

### Determination of HIV-1 infectious titer and bNAb concentration using TZM.bl assay

Determination of Tissue Culture Infectious Dose 50 (TCID50) of HIV-1 culture and plasma antibody concentration by neutralization assay was performed using a luciferase-based TZM.bl assay as previously described^20^. An equimolar tri-mix of 3BNC117, 10-1074 and SF12 was used for generating a standard curve for measuring tri-mix antibody levels from mouse plasma. The plasma was heated to 56°C for 30 mins prior to use in TZM.bl assay for inactivation of complement proteins. Bioluminescence was measured using Luciferin/lysis buffer composed of 10 mM MgCl2, 0.3 mM ATP, 0.5 mM Coenzyme A, 17 mM IGEPAL (all Sigma-Aldrich) and 1 mM D-Luciferin (GoldBio) in Tris-HCl on a BertholdTech TriStar2S luminometer.

### HIV-1 challenge of mice and viral load measurements

CD34 and CD34T+ mice were challenged on consecutive days intra-rectally with a total of 2.9 x 10^5^ TCID50 NL4-3_YU2_ by pipetting virus at the rectal opening without causing any injury in order to avoid direct systemic infection due to bleeding. Mice in the tri-mix group received a single intra-peritoneal injection of 2 mg each of 3BNC117, 10-1074 and SF12 in PBS, 24 hours prior to i.r. challenge. CD34 mice that remained un-infected after i.r. challenge, were challenged with HIV-1 via i.p. injection on consecutive days with a total of 7.3 x 10^5^ TCID50 NL4-3_YU2_. For viral load measurements, HIV-1 RNA was isolated using the QIAcube (Qiagen) from mouse plasma using the QIAamp MinElute Virus Spin Kit (Qiagen) along with a DNaseI (Qiagen) digestion step. Viral loads were determined by quantitative real-time PCR using *gag-*specific primers 6F 5’-CATGTTTTCAGCATTATCAGAAGGA-3’ and 84R 5’-TGCTTGATGTCCCCCCACT-3’ and *gag*-specific probe 56-FAM/CCACCCCACAAGATTTAAACACCATGCTAA /ZenDQ as previously described^21^. qPCR was performed on a Light Cycler 480 II (Roche) using the TaqMan RNA-to-CT 1-Step Kit (Thermo Fisher). An HIV-1 standard, whose copy number was determined using the Cobas 6800 HIV-1 kit (Roche) was included in every RNA isolation/qPCR run, was produced by super infection of SupT1-R5 cells. The limit of quantification of the qPCR was determined to be 384 HIV-1 RNA copies/ml and all values below this limit including mice that remained completely negative were assigned values between 100-300 for plotting graphs in GraphPad Prism 7.

### Production of bNAbs using 293-6E cells

3BNC117, 10-1074 and SF12 heavy and light chains were previously cloned into human antibody expression plasmids^15-17^. Antibodies were produced by transfection of 293-6E cells (National Research Council Canada) using branched polyethylenimine (PEI) 25kDa (Sigma-Aldrich). Cells were maintained at 37°C and 6% CO_2_ FreeStyle 293 Expression Medium (Thermo Fisher) and 0.2% Penicillin/Streptomycin 7 days post transfection, the cell culture supernatant was harvested, filtered with a 0.45 μM Nalgene Rapid Flow filter (Thermo Fisher) and incubated overnight at 4°C with Protein G Sepharose 4 Fast Flow (GE Healthcare) overnight. Antibody bound Sepharose beads were washed on chromatography columns (BioRad) and antibodies were eluted using 0.1M Glycine pH=3 and immediately buffered in 1M Tris pH=8. Thereafter, buffer exchange to PBS was performed using 50 kDa Amicon Ultra-15 spin columns (Millipore) and the antibodies were sterile filtered using 0.22 μm Ultrafree-CL columns (Millipore) and stored at 4°C.

### Histological characterization of mouse tissue

Mice were sacrificed using cervical dislocation and spleen, large intestine (colon) and small intestine (ileum) were excised. Mouse tissues were stored in 30% sucrose after overnight fixation with 4% PFA in PBS (Gibco) until freezing in Tissue freezing medium (Leica Biosytems). 10µm cryostat tissue sections were cut, mounted on SuperFrost Plus slides (Thermo Scientific), and stored at −80°C until further use. Fluorescence HIV-1 RNA *in situ* hybridisation was performed using the RNAscope Multiplex Fluorescent Detection Reagent Kit v2 and the RNAscope Probe-V-HIV1-CladeB probe (both ACD, bio-techne) according to the manufacturer’s instructions with minor modifications. Fixed-frozen tissue sections were baked at 60°C, re-fixed with 4% PFA in PBS, treated with hydrogen peroxide, and after antigen-retrieval by heating and protease digestion, probes were hybridised and signal was amplified. Bound probe was detected with Opal 570 fluorophore (Perkin Elmer). Subsequently, slides were incubated overnight with antibodies against human CD3 (CD3-12, Abcam) and CD4 (EPR6855, Abcam) at 4°C and bound antibodies were detected by fluorescently labelled anti-rat AlexaFluor 488 and anti-rabbit IgG AlexaFluor 647 antibodies (Thermo Fisher Scientific). Nuclei were counterstained with DAPI (Thermo Fisher Scientific) and slides were embedded in Fluoromount-G (Thermo Fisher Scientific). Immunohistochemistry (IHC) for human CD3, CD4, and CD8 was carried out as follows: Fixed frozen tissue sections were thawed, re-fixed with 4% PFA in PBS, and after antigen retrieval incubated with antibodies against human CD3 (CD3-12, Abcam), CD4 (EPR6855, Abcam), and CD8 (C8/144B, Agilent Dako). Bound primary antibodies were detected by fluorescently labelled anti-mouse IgG AlexaFluor 488, anti-rabbit IgG AlexaFluor 555, and anti-rat AlexaFluor 647 antibodies (Thermo Fisher Scientific). Nuclei were counterstained with DAPI (Thermo Fisher Scientific) and slides were embedded in Fluoromount-G (Thermo Fisher Scientific). Signal was visualised using an inverted Leica DMi8 epifluorescence microscope and final image processing was done using Adobe Illustrator^®^.

### Single genome amplification of HIV-1 *env* from mouse plasma

Plasma RNA was extracted using the QIAamp MinElute Virus Spin Kit (Qiagen) along with a DNaseI (Qiagen) digestion step using the QIAcube (Qiagen). cDNA was generated from plasma RNA using the primer YB383 5’-TTTTTTTTTTTTTTTTTTTTTTTTRAAGCAC-3’ as per the manufacturer’s protocol for the Superscript III Reverse Transcriptase (Thermo Fisher Scientific). cDNA was additionally treated with Ribonuclease H (Thermo Fisher) for 20 min at 37°C and stored at −80°C. The HIV-1_YU2_ *env* cDNA was amplified by nested PCR using dilutions that would yield <30% positive PCR reactions so that >80% reactions would yield a product derived from a single virus particle. 1^St^ PCR was performed using the primers YB383 5’-TTTTTTTTTTTTTTTTTTTTTTTTRAAGCAC -3’ and YB50 5’-GGCTTAGGCATCTCCTATGGCAGGAAGAA-3’ with the cycling conditions: 98°C for 45 s, 35 cycles of 98°C for 15 s, 55°C for 30 s and 72°C for 4 min and final amplification at 72°C for 15 min. 1 μl of the 1^st^ PCR product was used for 2^nd^ PCR with the primers YB49 5’-TAGAAAGAGCAGAAGACAGTGGCAATGA-3’ and YB52 5’-GGTGTGTAGTTCTGCCAATCA GGGAAGWAGCCTTGTG-3’ with the cycling conditions: 98°C for 45 s, 45 cycles of 98°C for 15 s, 55°C for 30 s and 72°C for 4 min and final amplification at 72°C for 15 min. PCR was performed using Phusion Hot Start Flex DNA Polymerase (New England Biolabs).

### Illumina dye sequencing of HIV-1 *env* amplicons and sequence analysis

NGS library preparation for Illumina Dye sequencing was done as previously described^22^. In brief, the 2^nd^ SGA *env* PCR products were cleaved into approximately 300 bp products by tagmentation using the Nextera DNA Library Prep Kit (Illumina). Indices from the Nextera Index Kit (Illumina), followed by adaptors P1 (AATGATACGGCGACCACCGA) and P2 (CAAGCAGAAGACGGCATACGA) were added by limited cycle PCR using the KAPA HiFi Hot Start Ready Mix (Roche). PCR products were purified using AMPure XP beads (Beckman Coulter), pooled and then sequenced using the MiSeq 300-cycle Nano Kit v2 (Illumina) spiked with ∽10% PhiX. Paired end reads were assembled as previously described^23^ and a consensus sequence was built. Further analysis was done using Geneious R10v10.0.9 where sequences with >75% nucleotide identity across reads were considered. Full-length sequences with high base quality and a maximum of 1 ambiguity were considered for final analysis. Alignments and phylogenetic trees were built using the ClustalOmega v1.2.3 and FastTree v2.1.11 plugins in Geneious R10v10.0.9.

