## Supplementary information for "A novel humanized mouse model to study mucosal HIV-1 transmission and prevention"

### Supplementary Figures


| | |
|---|---|
| Supplementary Figure 1 | *Analysis of leukocyte populations in the blood of CD34 and CD34T+ mice* |
| Supplementary Figure 2 | *Analysis of leukocyte populations in the tissue of CD34 and CD34T+ mice* |
| Supplementary Figure 3 | *Analysis of HIV-1 infection in CD34 mice* |
| Supplementary Figure 4 | *Analysis of SGS-derived env sequences from plasma of HIV-1 infected mice* |
| Supplementary Figure 5 | *Neutralization potency of bNAb tri-mix against NL4-3$_{YU2}$ HIV-1* |


### Supplementary Tables


| | |
|---|---|
| Supplementary Table 1 | *Leukocyte cell numbers in the blood of CD34T+ mice* |
| Supplementary Table 2 | *Leukocyte cell numbers in the blood of CD34 mice* |
| Supplementary Table 3 | *Leukocyte cell numbers in tissue of CD34T+ and CD34 mice* |

**a**

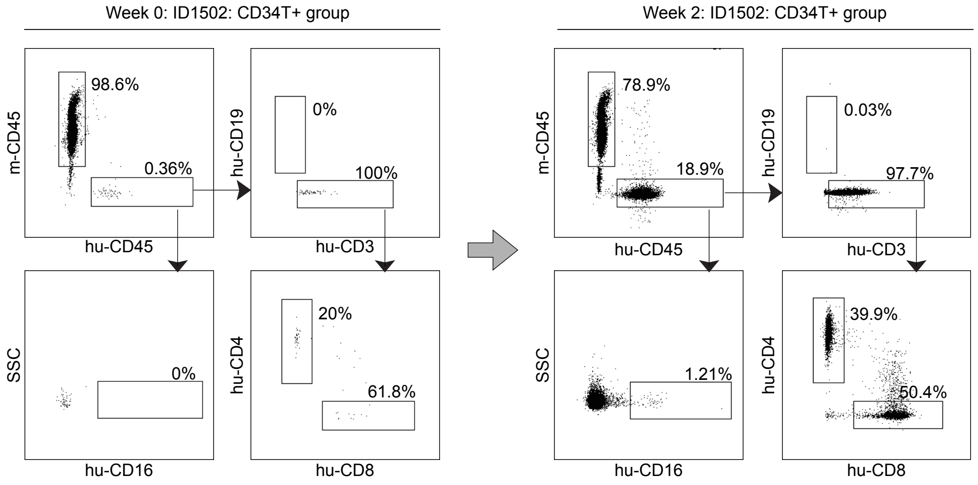


**b**

CD34 mice N=42

| Cell population | Mean cells/µl w0 | Mean cells/µl w2 | Fold expansion |
|---|---|---|---|
| hu-CD45+ | 95.9 | 83.0 | 0.9 |
| hu-CD3+ | 21.3 | 24.1 | 1.1 |
| hu-CD4+ | 11.9 | 15.3 | 1.3 |
| hu-CD8+ | 5.7 | 7.2 | 1.3 |
| hu-CD19+ | 67.8 | 51.5 | 0.8 |
| hu-CD16+ | 1.0 | 1.8 | 1.7 |
| mo-CD45+ | 818.1 | 827.7 | 1.0 |

CD34T+ mice N=69

| Cell population | Mean cells/µl w0 | Mean cells/µl w2 | Fold expansion |
|---|---|---|---|
| hu-CD45+ | 72.5 | 498.2 | 6.9 |
| hu-CD3+ | 23.2 | 448.9 | 19.3 |
| hu-CD4+ | 14.1 | 229.4 | 16.3 |
| hu-CD8+ | 7.3 | 190.5 | 26.2 |
| hu-CD19+ | 44.4 | 20.1 | 0.5 |
| hu-CD16+ | 0.9 | 20.4 | 23.6 |
| mo-CD45+ | 989.4 | 1243.6 | 1.3 |

| Fold expansion |
|---|
| <5 |
| 5-15 |
| >15 |

**c**

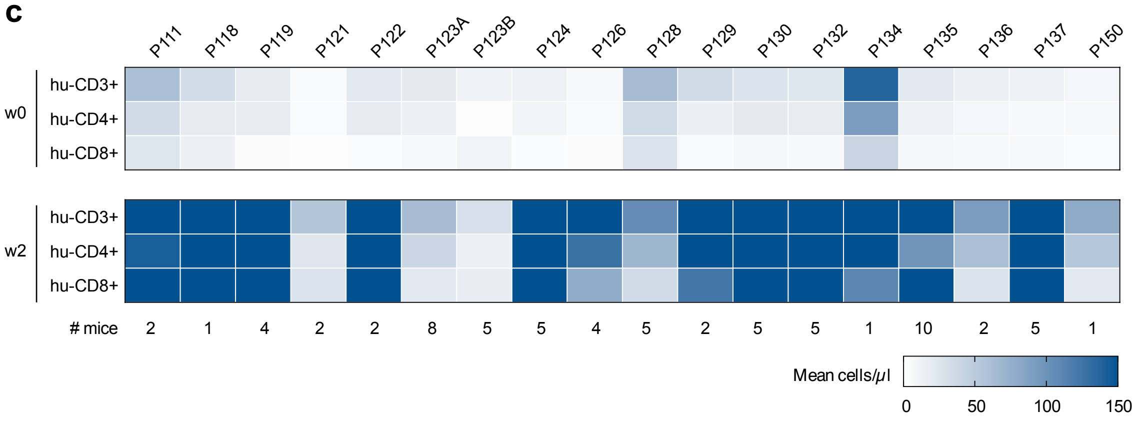


**Supplementary Figure 1: Analysis of leukocyte populations in the blood of CD34 and CD34T+ mice**
**a**, Gating strategy depicted with representative FACS plots from analysis of blood from a CD34T+ mouse ID1502 before (week 0) and after (week 2) UCBC and human IL-7 treatment. **b**, Mean cell numbers per µl blood and fold change between week 0 (w0) and week 2 (w2) from CD34 and CD34T+ mice shown in Fig. 1b. **c**, Analysis of human leukocyte expansion based on mean cell numbers per µl blood in CD34T+ mice (N=69) divided according to the cord blood donor.

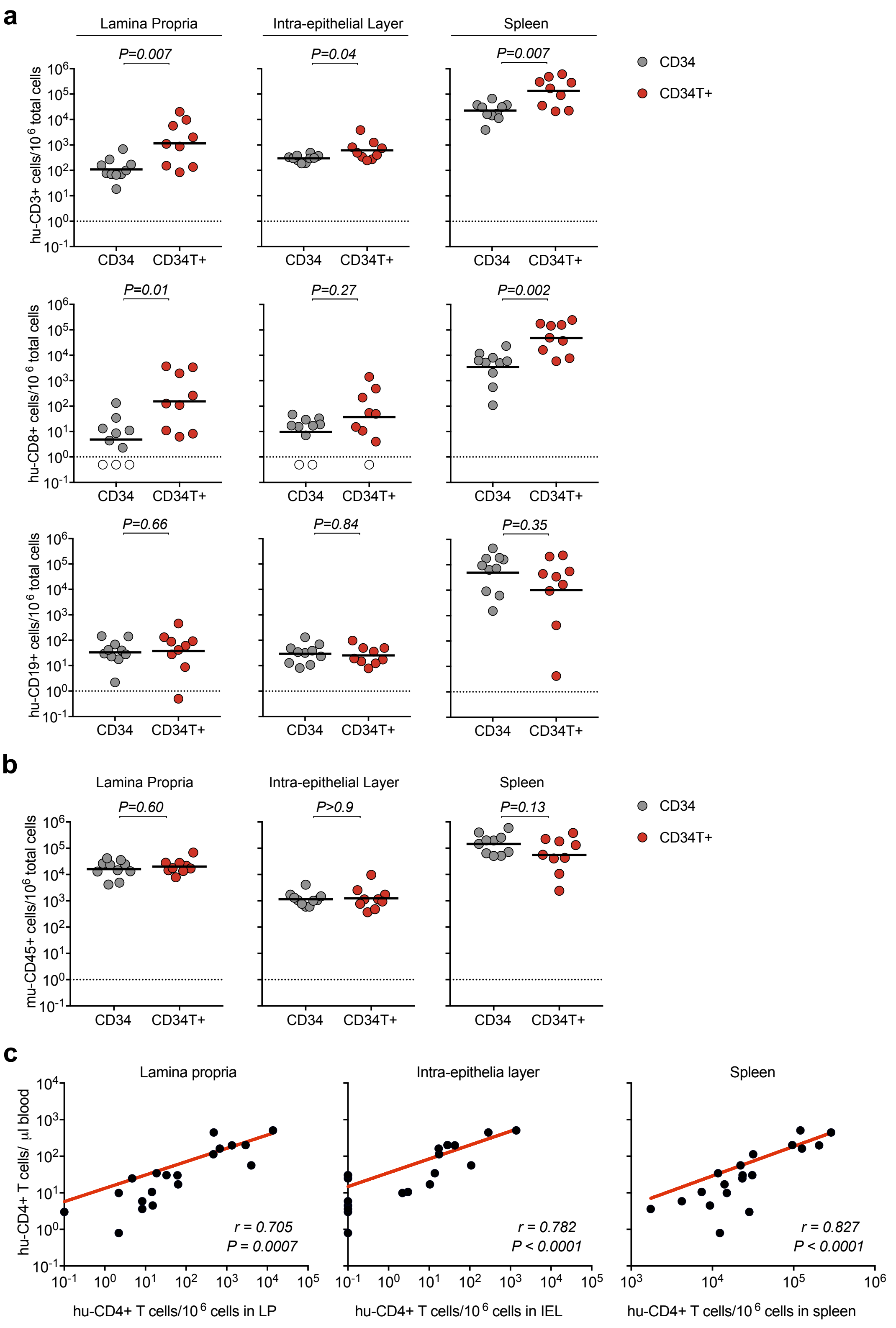


**Supplementary Figure 2: Analysis of leukocyte populations in the tissue of CD34 and CD34T+ mice**
FACS analysis of CD34 and CD34T+ mice shown in Fig. 2c for the presence of human CD3+ T cell, human CD8+ T cells, human CD19+ B cells (**a**) and mouse CD45+ cells (**b**) in the gut LP, IEL and the spleen at week 2. Points shown in white were below limit of detection of assay. Statistical analysis done using an Mann-Whitney U test. **c**, Combined Pearson correlation curves plotted for CD34 (N=10) and CD34T+(N=9) mice to analyze the relationship between human CD4+T cell levels in the blood vs. lymphoid tissue: LP, IEL and spleen).

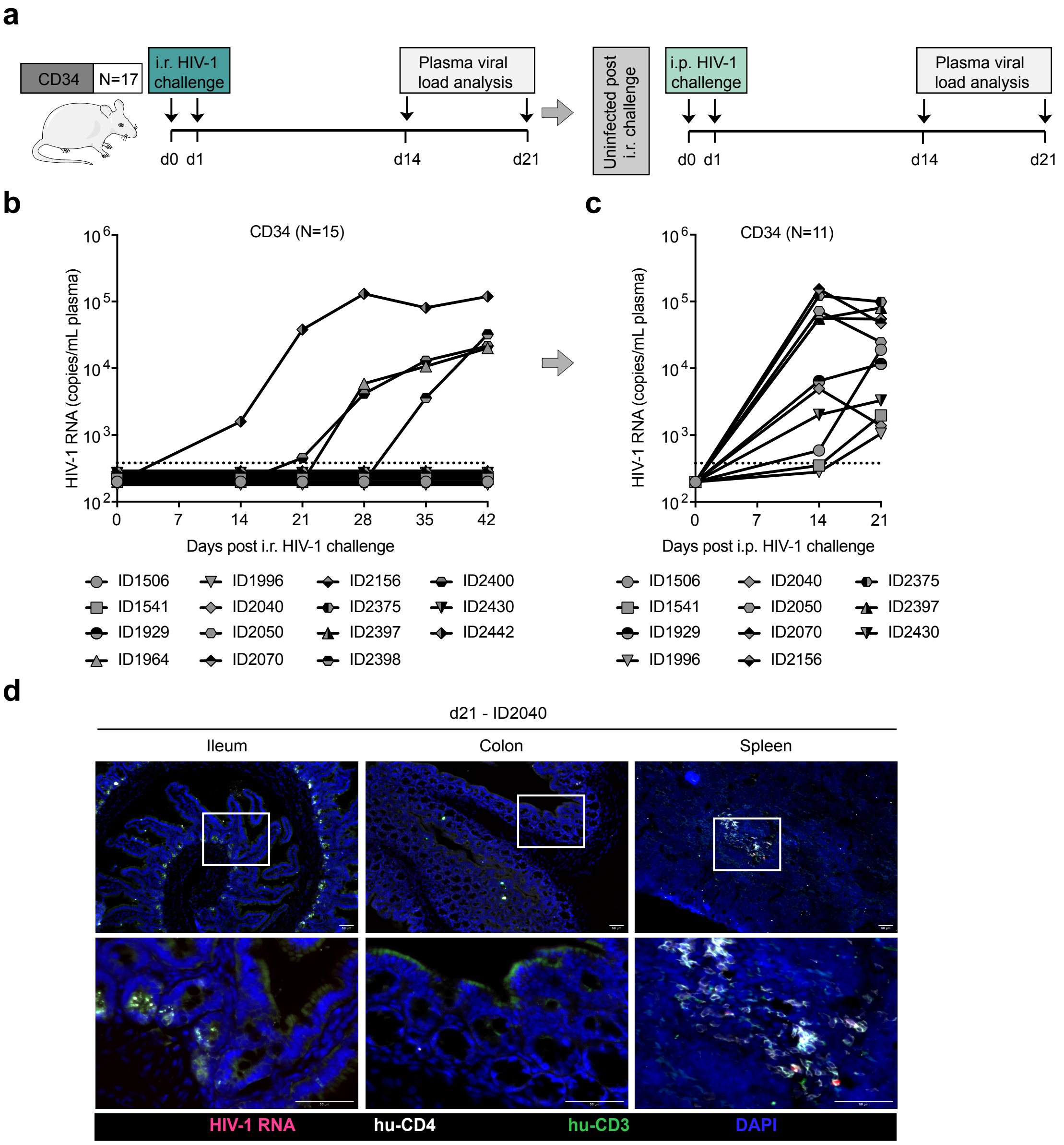


**Supplementary Figure 3:Supplementary Figure 3: Analysis of HIV-1 infection in CD34 mice**

**a**, Schematic representation of the experiment design. CD34 mice were given i.r. challenges with NL4-3YU2 HIV-1 on consecutive days and bled at the indicated time points for testing viremia. Mice that remained un-infected after i.r. challenge were challenged via i.p. injection of NL4-3YU2 HIV-1 to confirm susceptibility to infection. **b**, Plasma HIV-1 RNA levels in the blood of CD34 mice after i.r. challenge with high dose NL4-3YU2 HIV-1. **c**, Plasma HIV-1 RNA levels in CD34 mice after i.p. challenge with NL4-3YU2 HIV-1. The dotted line represents the detection limit of 384 HIV-1 RNA copies/ml of the assay in (b) and (c). **d**, RNA-scope detection of HIV-1 infection in the ileum, colon and spleen of CD34+ mice harvested at day 21 post i.p. HIV-1 challenge. HIV-1 RNA (pink), CD4+ (white), CD3+ (green) cells were labeled for detection along with DAPI for visualizing the nucleus. Scale bar is 50 μm. 3 mice per group were analyzed with 3 different slices from each mouse tissue being analyzed. Representative images were selected for display. i.r. intra-rectal, i.p. intra-peritoneal

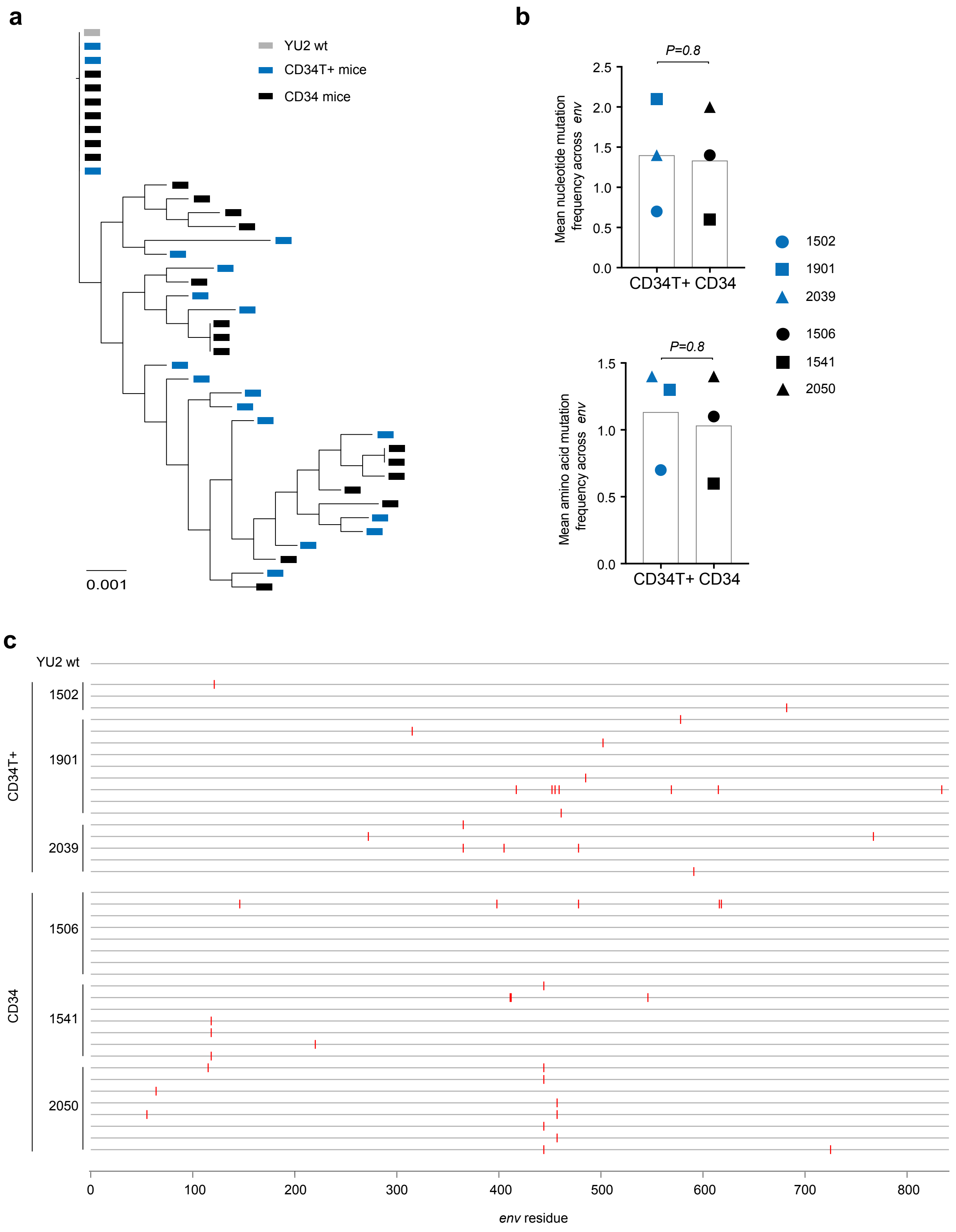


**Supplementary Figure 4: Analysis of SGS-derived *env* sequences from plasma of HIV-1 infected mice**
**a**, Phylogenetic tree of SGS-derived env sequences from the plasma of CD34T+ mice and CD34 mice at day 21 post post i.r. or i.p. challenge with NL4-3YU2 HIV-1 respectively. **b**, Analysis of mutation rates at the amino acid and nucleotide level in the env gene of mice from a. **c**, Amino acid alignment of plasma SGS-derived env sequences from mice analyzed in (**a** and **b**). Red bars indicate amino acid residues with mutations relative to the wild-type NL4-3YU2 HIV-1 env. Amino acid numbering is based on HIV-1YU2 env. i.r. intra-rectal, i.p. intra-peritoneal

**a**

| Epitope | bNAb | NL4-3YU2 HIV-1 IC50 | IC80 | MPI |
|---|---|---|---|---|
| V3 loop | 10-1074 | 0.812 | 2.426 | 100 |
| CD4bs | 3BNC117 | 0.052 | 0.138 | 100 |
| Silent face | SF12 | 0.914 | 2.456 | 100 |
| Combi. | Tri-mix | 0.107 | 0.320 | 100 |

| | | | | |
|---|---|---|---|---|
| IC50 (μg/ml) | < 0,5 | 0,5 - 2,5 | 2,5 - 10,0 | >10 |
| MPI | 90-100 | 50-90 | 25-50 | 0-25 |

**b**

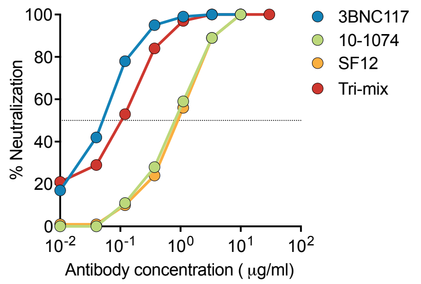


**c**

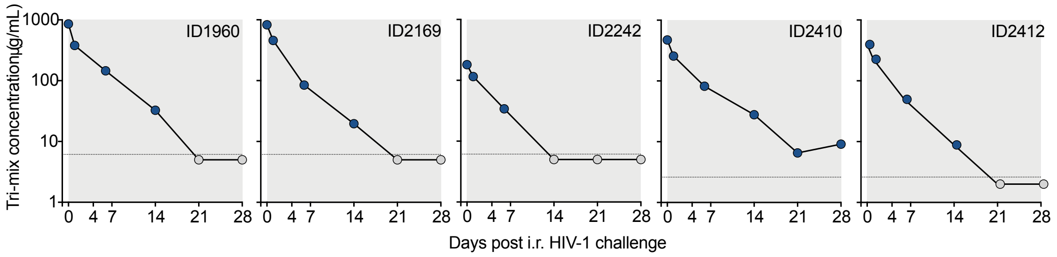


**d**

Plasma ID50 antibody levels against NL4-3 YU2

| TP | ID1960 | ID2242 | ID2169 | ID2410 | ID2412 |
|---|---|---|---|---|---|
| d0 | 6308 | 1330 | 6091 | 8155 | 6819 |
| d1 | 2788 | 849 | 3361 | 4432 | 3901 |
| d6 | 1068 | 250 | 622 | 1412 | 854 |
| d14 | 240 | 45 | 144 | 480 | 152 |
| d21 | <45 | <45 | <45 | 113 | <45 |
| d28 | <45 | <45 | <45 | 157 | <45 |
| d35 | <45 | <45 | <45 | <45 | <45 |
| d42 | <45 | <45 | <45 | 56 | <45 |
| d55 | <45 | <45 | <45 | 70 | <45 |

**e**

Plasma antibody levels in mice

| TP | ID1960 | ID2242 | ID2169 | ID2410 | ID2412 |
|---|---|---|---|---|---|
| d0 | 857.89 | 180.88 | 828.38 | 472.99 | 395.5 |
| d1 | 379.17 | 115.46 | 457.1 | 257.06 | 226.26 |
| d6 | 145.25 | 34 | 84.59 | 81.9 | 49.53 |
| d14 | 32.64 | 6.12 | 19.58 | 27.84 | 8.82 |
| d21 | 6.12 | 6.12 | 6.12 | 6.55 | 2.61 |
| d28 | 6.12 | 6.12 | 6.12 | 9.11 | 2.61 |
| d35 | 6.12 | 6.12 | 6.12 | 2.61 | 2.61 |
| d42 | 6.12 | 6.12 | 6.12 | 3.25 | 2.61 |
| d55 | 6.12 | 6.12 | 6.12 | 4.06 | 2.61 |

**Supplementary Figure 5: Neutralization potency of bNAb tri-mix against NL4-3YU2 HIV-1**
**a**, Neutralization potency of individual bNAbs (3BNC117, 10-1074 and SF12) or an equimolar tri-mix (combi.) against the NL4-3YU2 HIV-1 infectious molecular clone represented as 50% inhibitory concentration (IC50) or 80% inhibitory concentration (IC80) along with the maximum percent inhibition (MPI) values. **b**, Neutralization curve of the individual bNAbs along with the tri-mix measured in (a). **c**, Antibody concentration of the tri-mix in the plasma of CD34T+ tri-mix mice from Fig. 3 measured using the TZM-bl assay. **d**, 50% Inhibitory dilution (ID50) values of the tri-mix in the plasma of CD34T+ tri-mix mice from Fig. 3 measured against the NL4-3YU2 HIV-1 infectious molecular clone. ID50 values at the time of i.r. HIV-1 challenge (d0 and d1) are highlighted in red. **e**, Tri-mix concentration in the plasma of CD34T+ tri-mix mice from Fig. 3 measured against the NL4-3YU2 HIV-1 infectious molecular clone. Values at the time of i.r. HIV-1 challenge (d0 and d1) are highlighted in red and values below the detection limit of the assay are highlighted in grey.

**Supplementary Table 1.** Leukocyte cell numbers in the blood of CD34T+ mice

CD34T+ mice N=69

Number of cells/µl blood volume: <10 | 10-100 | >100

| Mouse ID | hu-CD45+ | | hu-CD3+ | | hu-CD4+ | | hu-CD8+ | | hu-CD19+ | | hu-CD16+ | | m-CD45+ | |
|---|---|---|---|---|---|---|---|---|---|---|---|---|---|---|
| | w0 | w2 | w0 | w2 | w0 | w2 | w0 | w2 | w0 | w2 | w0 | w2 | w0 | w2 |
| **1317** | 13,2 | 977,9 | 4,9 | 927,1 | 4,0 | 446,3 | 0,8 | 439,1 | 6,3 | 0,7 | nd | nd | 846,6 | 801,9 |
| **1320** | 41,3 | 1019,3 | 38,1 | 977,6 | 31,7 | 507,3 | 3,3 | 417,7 | 1,0 | 0,7 | nd | nd | 876,3 | 3291,4 |
| **1344** | 373,2 | 542,3 | 32,1 | 472,4 | 18,0 | 198,2 | 12,7 | 204,5 | 323,6 | 11,0 | nd | nd | 793,1 | 45,1 |
| **1550** | 98,1 | 632,7 | 54,8 | 615,6 | 32,7 | 202,0 | 19,1 | 354,6 | 38,4 | 9,2 | nd | nd | 719,4 | 201,3 |
| **1571** | 4,0 | 229,4 | 3,8 | 225,7 | 2,9 | 162,1 | 0,4 | 54,2 | 0,1 | 0,5 | nd | nd | 652,6 | 981,2 |
| **1618** | 0,9 | 22,2 | 0,7 | 21,5 | 0,1 | 9,9 | 0,5 | 8,8 | 0,0 | 0,1 | nd | nd | 673,3 | 911,8 |
| **1800** | 65,7 | 181,0 | 23,1 | 170,5 | 9,7 | 56,1 | 3,8 | 88,8 | 33,2 | 5,4 | nd | nd | 524,1 | 24,0 |
| **1551** | 78,5 | 260,0 | 62,9 | 256,2 | 31,9 | 79,8 | 27,1 | 156,8 | 10,6 | 1,7 | nd | nd | 538,6 | 219,3 |
| **1578** | 5,7 | 87,0 | 4,7 | 85,2 | 1,2 | 33,7 | 2,7 | 44,1 | 0,3 | 0,3 | nd | nd | 1874,5 | 1078,5 |
| **1614** | 32,2 | 140,4 | 31,9 | 139,1 | 26,3 | 87,9 | 4,2 | 45,4 | 0,0 | 0,0 | nd | nd | 1329,2 | 2288,0 |
| **1782** | 46,0 | 490,5 | 44,2 | 486,2 | 18,1 | 244,5 | 3,5 | 155,7 | 0,7 | 0,2 | nd | nd | 842,1 | 1930,3 |
| **1502** | 3,8 | 348,7 | 3,8 | 340,5 | 2,4 | 171,5 | 0,8 | 136,0 | 0,0 | 0,1 | 0,0 | 4,2 | 1052,2 | 1454,1 |
| **1508** | 63,8 | 141,0 | 63,2 | 139,7 | 59,4 | 100,0 | 2,9 | 32,7 | 0,0 | 0,1 | 0,1 | 1,2 | 1550,3 | 1322,3 |
| **1529** | 2,9 | 135,8 | 2,8 | 131,6 | 0,6 | 43,1 | 1,9 | 72,2 | 0,0 | 0,1 | 0,0 | 1,8 | 670,3 | 865,0 |
| **1534** | 2,1 | 1039,1 | 2,1 | 1014,1 | 1,4 | 406,3 | 0,7 | 470,8 | 0,0 | 0,4 | 0,0 | 23,0 | 1101,6 | 786,0 |
| **1539** | 0,5 | 260,9 | 0,4 | 256,1 | 0,1 | 135,3 | 0,2 | 109,3 | 0,0 | 0,0 | 0,0 | 2,8 | 510,3 | 2004,4 |
| **1542** | 12,5 | 289,6 | 12,5 | 282,8 | 8,8 | 174,9 | 3,5 | 99,1 | 0,0 | 0,2 | 0,0 | 4,4 | 1239,9 | 2861,7 |
| **1901** | 0,1 | 39,1 | 0,1 | 16,3 | 0,0 | 7,1 | 0,1 | 7,1 | 0,0 | 0,1 | 0,0 | 17,4 | 462,7 | 1184,8 |
| **1915** | 7,5 | 134,9 | 7,5 | 89,9 | 6,0 | 38,8 | 1,5 | 44,4 | 0,0 | 0,2 | 0,0 | 32,0 | 999,0 | 1774,1 |
| **2021** | 37,5 | 73,2 | 11,6 | 56,8 | 8,6 | 29,4 | 2,5 | 21,8 | 18,5 | 11,0 | 0,2 | 1,2 | 373,5 | 312,8 |
| **2039** | 13,2 | 97,2 | 0,6 | 94,2 | 0,3 | 33,8 | 0,1 | 44,7 | 10,6 | 0,4 | 0,1 | 1,8 | 553,4 | 418,4 |
| **2047** | 15,9 | 2065,4 | 9,6 | 2026,4 | 6,4 | 1197,6 | 2,1 | 723,4 | 1,8 | 0,0 | 0,9 | 32,2 | 301,3 | 1075,9 |
| **1930** | 16,8 | 64,7 | 6,5 | 23,1 | 5,4 | 17,6 | 0,9 | 3,6 | 7,4 | 29,7 | 0,3 | 5,3 | 342,3 | 1122,0 |
| **1931** | 50,8 | 164,6 | 29,7 | 117,9 | 22,3 | 81,3 | 6,3 | 31,8 | 14,1 | 35,6 | 2,2 | 7,1 | 547,8 | 764,7 |
| **1956** | 43,5 | 41,8 | 4,8 | 8,0 | 2,2 | 3,8 | 2,1 | 3,7 | 33,8 | 21,6 | 0,4 | 1,0 | 2004,9 | 10542 |
| **1958** | 36,9 | 44,8 | 10,1 | 19,7 | 7,3 | 13,4 | 2,2 | 5,0 | 22,4 | 19,9 | 0,8 | 2,3 | 1369,7 | 808,1 |
| **1959** | 50,0 | 127,2 | 12,8 | 45,8 | 9,2 | 32,7 | 2,5 | 10,8 | 29,3 | 54,7 | 3,7 | 20,1 | 657,2 | 812,5 |
| **1960** | 29,9 | 49,6 | 16,5 | 28,2 | 11,9 | 17,8 | 3,4 | 7,3 | 9,9 | 15,8 | 1,1 | 2,5 | 654,0 | 589,1 |
| **1961** | 60,8 | 139,2 | 36,4 | 121,3 | 19,8 | 65,3 | 14,6 | 48,6 | 14,6 | 7,3 | 5,2 | 10,1 | 287,3 | 95,6 |
| **1962** | 85,2 | 210,1 | 30,7 | 128,2 | 23,8 | 70,5 | 5,2 | 48,7 | 43,1 | 58,7 | 2,7 | 17,0 | 320,0 | 226,2 |
| **2071** | 43,7 | 43,3 | 42,8 | 42,4 | 0,1 | 13,8 | 39,6 | 26,4 | 0,0 | 0,0 | 0,1 | 0,1 | 3089,6 | 2544,1 |
| **2098** | 1,6 | 6,1 | 0,9 | 4,4 | 0,4 | 1,9 | 0,4 | 2,2 | 0,4 | 0,8 | 0,0 | 0,2 | 1153,0 | 1186,1 |
| **2099** | 0,9 | 15,3 | 0,9 | 14,8 | 0,9 | 10,4 | 0,0 | 4,1 | 0,0 | 0,0 | 0,0 | 0,1 | 1120,4 | 1237,4 |
| **2109** | 3,7 | 73,6 | 3,7 | 73,0 | 1,0 | 28,3 | 2,7 | 42,0 | 0,0 | 0,2 | 0,0 | 0,1 | 3764,3 | 5449,2 |
| **2149** | 102,4 | 288,0 | 32,2 | 204,6 | 16,0 | 105,7 | 15,3 | 88,7 | 57,7 | 63,6 | 1,3 | 7,5 | 702,7 | 690,1 |
| **2150** | 65,7 | 50,9 | 61,1 | 48,8 | 39,5 | 28,2 | 18,4 | 19,4 | 3,3 | 0,7 | 0,1 | 0,1 | 422,8 | 397,1 |
| **2154** | 258,6 | 199,6 | 176,1 | 179,2 | 90,3 | 149,9 | 82,1 | 26,7 | 73,3 | 16,6 | 1,2 | 0,5 | 986,5 | 768,7 |
| **2168** | 70,9 | 132,8 | 11,1 | 50,5 | 7,9 | 32,7 | 2,7 | 14,7 | 52,7 | 55,9 | 0,9 | 11,2 | 472,2 | 684,0 |
| **2169** | 31,6 | 37,5 | 30,8 | 37,0 | 18,6 | 20,6 | 11,3 | 14,7 | 0,2 | 0,1 | 0,1 | 0,1 | 528,0 | 797,8 |
| **2242** | 1,2 | 10,9 | 1,1 | 8,0 | 0,9 | 6,5 | 0,1 | 0,9 | 0,0 | 2,7 | 0,0 | 0,1 | 938,9 | 2610,6 |
| **2372** | 28,7 | 62,4 | 7,6 | 23,8 | 6,0 | 17,1 | 1,4 | 6,1 | 17,7 | 32,6 | 0,4 | 1,7 | 560,2 | 877,8 |
| **2381** | 211,7 | 305,5 | 3,8 | 54,2 | 1,2 | 34,4 | 2,5 | 11,0 | 191,6 | 201,5 | 1,4 | 29,5 | 1054,7 | 619,5 |
| **2373** | 51,2 | 1924,0 | 48,7 | 1906,8 | 30,2 | 1159,6 | 15,3 | 684,5 | 1,4 | 0,4 | 0,5 | 2,0 | 810,4 | 1742,6 |
| **2376** | 61,5 | 513,5 | 17,2 | 499,4 | 11,9 | 273,9 | 3,4 | 199,9 | 36,2 | 1,1 | 1,1 | 8,9 | 1263,2 | 542,6 |
| **2405** | 282,0 | 444,6 | 49,9 | 278,5 | 41,7 | 167,0 | 7,1 | 105,8 | 211,8 | 127,1 | 3,7 | 20,4 | 610,9 | 249,3 |
| **2410** | 55,2 | 146,0 | 54,1 | 143,0 | 41,1 | 35,8 | 8,6 | 101,6 | 0,9 | 0,0 | 0,0 | 0,2 | 1237,9 | 1215,2 |
| **2412** | 69,5 | 155,1 | 17,8 | 146,1 | 5,6 | 78,7 | 10,4 | 59,6 | 44,9 | 2,5 | 3,4 | 1,2 | 1307,0 | 963,7 |
| **2415** | 88,7 | 1101,7 | 81,3 | 1081,6 | 65,7 | 741,4 | 10,3 | 306,1 | 5,6 | 0,0 | 0,1 | 14,8 | 1907,1 | 607,1 |
| **2417** | 81,0 | 1762,5 | 9,5 | 1634,3 | 7,4 | 949,5 | 1,8 | 595,1 | 63,9 | 5,7 | 0,7 | 69,9 | 876,5 | 551,8 |
| **2418** | 159,1 | 4613,0 | 9,6 | 4093,2 | 7,0 | 2064,8 | 2,5 | 1769,1 | 136,3 | 29,8 | 0,5 | 338,3 | 1766,9 | 1445,9 |
| **2419** | 200,7 | 939,5 | 7,0 | 597,8 | 2,3 | 180,7 | 4,3 | 372,7 | 172,1 | 45,3 | 1,0 | 232,0 | 804,3 | 26,4 |
| **2422** | 79,3 | 1621,1 | 5,2 | 1489,8 | 2,5 | 892,5 | 2,6 | 476,9 | 70,5 | 3,2 | 0,7 | 91,0 | 1153,6 | 372,7 |
| **2424** | 52,1 | 238,4 | 19,2 | 176,6 | 13,8 | 73,2 | 5,2 | 85,3 | 30,4 | 44,3 | 0,7 | 14,4 | 1743,1 | 620,6 |
| **2425** | 32,6 | 301,6 | 5,4 | 281,9 | 3,4 | 95,4 | 1,3 | 157,8 | 24,9 | 5,6 | 0,2 | 8,7 | 882,2 | 949,5 |
| **2426** | 37,7 | 339,9 | 14,5 | 336,0 | 12,2 | 171,0 | 1,1 | 146,1 | 22,1 | 0,3 | 0,1 | 0,6 | 975,7 | 1846,7 |
| **2429** | 51,5 | 209,2 | 6,3 | 173,3 | 4,1 | 75,3 | 2,0 | 76,8 | 39,1 | 24,0 | 0,3 | 5,6 | 611,9 | 776,6 |
| **2431** | 91,2 | 787,3 | 23,6 | 755,1 | 10,5 | 244,8 | 9,6 | 462,9 | 58,5 | 4,4 | 1,2 | 14,1 | 1010,1 | 824,5 |
| **2433** | 285,0 | 384,4 | 6,7 | 246,8 | 1,0 | 75,0 | 2,8 | 138,3 | 262,1 | 75,5 | 2,2 | 49,7 | 722,5 | 94,5 |
| **2434** | 563,8 | 409,6 | 58,8 | 235,7 | 20,9 | 62,3 | 27,2 | 157,4 | 482,9 | 154,4 | 4,1 | 11,5 | 541,4 | 45,9 |
| **2435** | 60,8 | 322,2 | 5,4 | 263,0 | 3,1 | 64,7 | 2,0 | 182,5 | 50,2 | 49,2 | 0,4 | 6,0 | 1184,5 | 156,9 |
| **2436** | 35,8 | 230,0 | 2,9 | 221,1 | 1,6 | 53,9 | 1,0 | 128,7 | 29,9 | 3,5 | 0,2 | 1,7 | 1071,0 | 1344,2 |
| **2438** | 22,3 | 437,5 | 2,6 | 419,5 | 1,0 | 213,5 | 1,5 | 169,0 | 18,2 | 2,7 | 0,2 | 5,5 | 854,5 | 1775,7 |
| **2439** | 12,0 | 215,1 | 4,0 | 200,6 | 2,0 | 117,7 | 1,8 | 63,4 | 6,7 | 0,5 | 0,1 | 7,6 | 1854,1 | 993,7 |
| **2443** | 94,0 | 251,2 | 5,1 | 205,9 | 2,7 | 84,0 | 2,4 | 103,8 | 85,6 | 24,8 | 0,7 | 6,8 | 1361,4 | 841,0 |
| **2444** | 133,9 | 4889,2 | 44,0 | 4759,8 | 16,9 | 2260,5 | 16,9 | 2296,9 | 86,5 | 70,1 | 1,1 | 8,4 | 959,4 | 676,9 |
| **2551** | 118,1 | 172,1 | 7,4 | 78,1 | 4,3 | 52,1 | 2,9 | 20,2 | 104,8 | 71,7 | 1,1 | 15,8 | 914,2 | 672,9 |
| **2571** | 18,6 | 160,0 | 15,2 | 149,3 | 11,4 | 100,2 | 3,7 | 40,4 | 0,0 | 0,7 | 2,4 | 8,2 | 432,4 | 1202,3 |
| **2575** | 10,3 | 31,4 | 9,9 | 30,5 | 3,4 | 16,6 | 5,9 | 11,1 | 0,0 | 0,1 | 0,0 | 0,1 | 877,1 | 6681,7 |
| **2649** | 142,4 | 591,6 | 136,7 | 578,6 | 89,8 | 446,8 | 40,0 | 108,9 | 2,4 | 3,2 | 0,2 | 6,0 | 1077,0 | 1671,3 |

**Supplementary Table 2.** Leukocyte cell numbers in the blood of CD34 mice

CD34 mice N=42

Number of cells/µl blood volume: <10 | 10-100 | >100

| Mouse ID | hu-CD45+ w0 | hu-CD45+ w2 | hu-CD3+ w0 | hu-CD3+ w2 | hu-CD4+ w0 | hu-CD4+ w2 | hu-CD8+ w0 | hu-CD8+ w2 | hu-CD19+ w0 | hu-CD19+ w2 | hu-CD16+ w0 | hu-CD16+ w2 | mu-CD45+ w0 | mu-CD45+ w2 |
|---|---|---|---|---|---|---|---|---|---|---|---|---|---|---|
| **1300** | 447,3 | 258,2 | 4,5 | 7,1 | 12,2 | 24,8 | 17,4 | 33,5 | 408,2 | 193,4 | nd | nd | 641,4 | 255,0 |
| **1306** | 493,7 | 235,6 | 6,6 | 6,9 | 14,2 | 10,6 | 23,3 | 18,5 | 456,9 | 209,5 | nd | nd | 524,2 | 195,2 |
| **1321** | 42,2 | 134,3 | 5,7 | 10,6 | 24,2 | 113,0 | 31,0 | 124,9 | 9,9 | 0,4 | nd | nd | 803,2 | 773,2 |
| **1337** | 4,2 | 3,9 | 0,5 | 0,1 | 0,8 | 0,8 | 1,5 | 0,9 | 1,7 | 0,9 | nd | nd | 2340,8 | 3357,8 |
| **1548** | 45,2 | 54,6 | 23,8 | 21,5 | 19,3 | 30,3 | 44,9 | 54,4 | 0,1 | 0,0 | nd | nd | 804,5 | 591,7 |
| **1576** | 60,7 | 86,5 | 4,7 | 6,7 | 14,1 | 30,0 | 19,4 | 37,3 | 33,9 | 38,2 | nd | nd | 899,1 | 694,7 |
| **1620** | 7,0 | 17,1 | 0,0 | 0,6 | 1,6 | 3,0 | 1,7 | 3,9 | 4,5 | 11,5 | nd | nd | 945,5 | 936,8 |
| **1802** | 26,8 | 64,9 | 2,4 | 2,3 | 2,6 | 4,5 | 5,4 | 7,7 | 16,3 | 49,9 | nd | nd | 485,3 | 619,3 |
| **1549** | 115,6 | 161,9 | 14,8 | 24,9 | 27,0 | 50,3 | 43,7 | 78,4 | 64,6 | 71,6 | nd | nd | 427,0 | 382,9 |
| **1617** | 21,3 | 9,8 | 2,2 | 1,1 | 15,5 | 6,7 | 18,9 | 8,6 | 1,0 | 0,7 | nd | nd | 1809,4 | 840,0 |
| **1619** | 0,3 | 0,6 | 0,1 | 0,3 | 0,1 | 0,2 | 0,3 | 0,6 | 0,0 | 0,0 | nd | nd | 1300,3 | 1041,1 |
| **1803** | 213,3 | 38,6 | 36,9 | 9,4 | 57,5 | 13,9 | 210,2 | 38,2 | 0,7 | 0,1 | nd | nd | 381,1 | 1096,2 |
| **1506** | 1,9 | 1,5 | 0,4 | 0,5 | 1,3 | 0,6 | 1,8 | 1,4 | 0,0 | 0,0 | 0,0 | 0,1 | 956,8 | 935,7 |
| **1535** | 18,8 | 0,3 | 0,3 | 0,1 | 0,2 | 0,1 | 0,7 | 0,3 | 15,1 | 0,0 | 0,1 | 0,0 | 466,9 | 338,9 |
| **1541** | 3,3 | 2,1 | 1,5 | 0,9 | 1,2 | 0,8 | 3,3 | 1,8 | 0,0 | 0,0 | 0,0 | 0,0 | 838,2 | 648,9 |
| **1996** | 14,2 | 18,4 | 1,9 | 1,6 | 1,5 | 2,7 | 3,7 | 4,8 | 7,8 | 10,6 | 0,0 | 0,0 | 510,1 | 636,7 |
| **2035** | 18,6 | 7,3 | 3,0 | 0,7 | 5,9 | 0,9 | 8,9 | 1,7 | 7,5 | 4,8 | 0,0 | 0,0 | 1004,8 | 547,8 |
| **2040** | 23,2 | 15,2 | 0,4 | 2,7 | 13,4 | 11,1 | 14,7 | 14,6 | 7,6 | 0,5 | 0,1 | 0,1 | 247,8 | 695,7 |
| **2044** | 1,7 | 2,4 | 0,1 | 0,2 | 0,3 | 0,2 | 0,4 | 0,5 | 0,9 | 1,0 | 0,0 | 0,1 | 544,6 | 1150,6 |
| **2048** | 5,6 | 2,2 | 1,9 | 0,7 | 3,5 | 1,2 | 5,6 | 2,1 | 0,0 | 0,0 | 0,0 | 0,1 | 875,3 | 1518,0 |
| **2049** | 74,9 | 20,0 | 4,4 | 2,8 | 63,8 | 16,1 | 74,5 | 19,7 | 0,1 | 0,0 | 0,1 | 0,1 | 617,4 | 1724,6 |
| **2050** | 17,3 | 47,0 | 0,1 | 0,5 | 2,6 | 19,3 | 3,2 | 20,4 | 8,3 | 19,1 | 0,6 | 1,2 | 274,4 | 164,9 |
| **1929** | 78,5 | 120,0 | 10,2 | 10,6 | 16,6 | 32,7 | 29,0 | 45,3 | 37,3 | 63,8 | 1,6 | 5,2 | 400,2 | 416,9 |
| **1964** | 91,6 | 115,3 | 9,4 | 12,8 | 28,7 | 27,1 | 40,3 | 42,8 | 33,3 | 54,5 | 8,4 | 8,5 | 480,0 | 477,4 |
| **2070** | 23,0 | 16,8 | 0,2 | 0,0 | 22,6 | 16,4 | 22,9 | 16,5 | 0,0 | 0,0 | 0,0 | 0,0 | 1043,7 | 1917,9 |
| **2151** | 86,8 | 266,9 | 26,0 | 104,0 | 34,9 | 99,0 | 65,5 | 217,6 | 8,5 | 21,4 | 1,8 | 10,5 | 617,1 | 789,3 |
| **2156** | 12,1 | 11,0 | 3,2 | 3,8 | 8,4 | 6,3 | 12,1 | 10,8 | 0,0 | 0,1 | 0,0 | 0,1 | 983,1 | 814,6 |
| **2171** | 47,8 | 89,2 | 10,0 | 22,4 | 9,3 | 27,2 | 19,7 | 51,6 | 21,2 | 21,2 | 0,6 | 3,2 | 215,4 | 261,1 |
| **2371** | 54,6 | 60,6 | 2,2 | 2,1 | 3,4 | 3,6 | 6,3 | 5,9 | 42,7 | 48,8 | 0,3 | 0,3 | 1123,7 | 852,4 |
| **2382** | 115,0 | 112,9 | 5,1 | 7,6 | 2,9 | 5,9 | 9,2 | 14,6 | 90,6 | 80,0 | 1,4 | 2,3 | 612,7 | 473,3 |
| **2375** | 158,3 | 133,2 | 1,5 | 1,8 | 15,1 | 7,6 | 17,6 | 10,4 | 127,6 | 110,0 | 2,0 | 2,0 | 1263,5 | 742,7 |
| **2378** | 513,2 | 340,8 | 1,4 | 0,8 | 8,0 | 5,0 | 9,8 | 6,5 | 468,2 | 311,6 | 5,8 | 6,9 | 521,0 | 329,6 |
| **2397** | 297,6 | 308,7 | 3,9 | 3,9 | 3,7 | 6,0 | 9,0 | 11,2 | 264,7 | 272,6 | 1,7 | 4,5 | 864,1 | 858,5 |
| **2398** | 132,0 | 208,5 | 3,1 | 3,0 | 5,8 | 13,3 | 10,3 | 17,5 | 111,6 | 171,7 | 0,6 | 3,8 | 1215,0 | 876,2 |
| **2400** | 247,1 | 284,7 | 4,2 | 2,2 | 4,1 | 15,7 | 9,4 | 19,4 | 213,9 | 237,1 | 2,0 | 4,5 | 1180,1 | 693,8 |
| **2421** | 50,4 | 26,8 | 10,4 | 11,4 | 8,5 | 9,5 | 21,1 | 22,9 | 26,2 | 2,0 | 1,2 | 0,1 | 695,1 | 1313,5 |
| **2430** | 70,4 | 72,1 | 7,8 | 4,1 | 5,9 | 6,4 | 13,7 | 10,9 | 50,8 | 53,0 | 0,3 | 1,0 | 610,8 | 986,5 |
| **2442** | 234,5 | 33,9 | 11,1 | 6,6 | 14,8 | 10,0 | 27,0 | 18,8 | 189,3 | 13,3 | 1,4 | 0,3 | 739,4 | 1138,2 |
| **2549** | 47,4 | 31,2 | 2,0 | 2,1 | 11,2 | 10,1 | 13,6 | 12,2 | 29,6 | 17,8 | 0,3 | 0,1 | 556,2 | 660,9 |
| **2550** | 84,7 | 84,6 | 0,3 | 1,0 | 0,6 | 0,4 | 1,2 | 1,8 | 78,6 | 79,1 | 0,9 | 1,0 | 1022,1 | 665,6 |
| **2573** | 3,9 | 4,5 | 1,7 | 2,2 | 1,0 | 1,5 | 2,8 | 4,3 | 0,6 | 0,0 | 0,0 | 0,0 | 1083,7 | 1105,9 |
| **2647** | 22,4 | 18,2 | 8,3 | 5,2 | 10,3 | 10,4 | 20,2 | 16,5 | 1,2 | 1,0 | 0,2 | 0,1 | 1433,4 | 501,7 |

**Supplementary Table 3.** Leukocyte cell numbers in tissue of CD34T+ and CD34 mice

| | | No. of cells/ $10^6$ cells in lamina propria at w2 | | | | |
|---|---|---|---|---|---|---|
| Model | Mouse ID | hu-CD3+ | hu-CD4+ | hu-CD8+ | hu-CD19+ | mu-CD45+ |
| CD34 | 1306 | 267.6 | 14.4 | 34.5 | 141.0 | 12975.0 |
| CD34 | 1321 | 688.6 | 468.6 | 131.5 | 145.8 | 4915.8 |
| CD34 | 1300 | 70.0 | 4.7 | 0.0 | 39.7 | 4167.8 |
| CD34 | 1337 | 99.3 | 2.2 | 11.0 | 17.7 | 24507.5 |
| CD34 | 1576 | 171.0 | 61.4 | 13.2 | 67.9 | 26189.6 |
| CD34 | 1802 | 77.3 | 15.0 | 8.6 | 27.9 | 23331.0 |
| CD34 | 1548 | 70.3 | 33.0 | 4.4 | 2.2 | 35584.3 |
| CD34 | 1620 | 18.3 | 0.0 | 2.3 | 29.8 | 14469.8 |
| CD34 | 2371 | 159.4 | 8.3 | 0.0 | 22.8 | 41694.4 |
| CD34 | 2382 | 66.8 | 8.3 | 0.0 | 41.7 | 13228.6 |
| CD34T+ | 1344 | 2003.4 | 1343.6 | 260.0 | 91.8 | 14545.6 |
| CD34T+ | 1317 | 868.3 | 477.4 | 126.2 | 131.1 | 7876.1 |
| CD34T+ | 1320 | 20039.9 | 13773.5 | 3346.8 | 453.1 | 68751.0 |
| CD34T+ | 1550 | 5709.2 | 2911.0 | 1937.1 | 60.7 | 21496.3 |
| CD34T+ | 1800 | 9681.3 | 4027.8 | 3662.0 | 27.8 | 17683.4 |
| CD34T+ | 1571 | 1099.5 | 675.1 | 109.9 | 88.0 | 27717.9 |
| CD34T+ | 1618 | 83.9 | 2.2 | 11.0 | 8.8 | 16987.1 |
| CD34T+ | 2372 | 135.7 | 63.7 | 8.2 | 0.0 | 28134.8 |
| CD34T+ | 2381 | 151.8 | 18.7 | 6.2 | 41.6 | 13640.2 |

| | | No. of cells/ $10^6$ cells in intra-epithelium at w2 | | | | |
|---|---|---|---|---|---|---|
| Model | Mouse ID | hu-CD3+ | hu-CD4+ | hu-CD8+ | hu-CD19+ | mu-CD45+ |
| CD34 | 1306 | 330.6 | 3.0 | 33.4 | 48.5 | 1052.5 |
| CD34 | 1321 | 250.5 | 17.6 | 7.1 | 31.7 | 1710.9 |
| CD34 | 1300 | 192.0 | 0.0 | 15.6 | 131.7 | 1049.5 |
| CD34 | 1337 | 372.1 | 0.0 | 29.3 | 69.9 | 4124.7 |
| CD34 | 1576 | 495.3 | 0.0 | 47.0 | 34.2 | 602.0 |
| CD34 | 1802 | 296.8 | 0.0 | 17.2 | 38.7 | 600.0 |
| CD34 | 1548 | 293.1 | 0.0 | 19.3 | 12.8 | 1320.2 |
| CD34 | 1620 | 186.0 | 0.0 | 17.1 | 10.7 | 1490.0 |
| CD34 | 2371 | 386.3 | 0.0 | 0.0 | 23.3 | 1001.9 |
| CD34 | 2382 | 307.2 | 0.0 | 0.0 | 8.1 | 767.9 |
| CD34T+ | 1344 | 271.1 | 42.9 | 54.2 | 49.9 | 368.7 |
| CD34T+ | 1317 | 1249.4 | 286.5 | 217.0 | 35.8 | 1727.7 |
| CD34T+ | 1320 | 3835.9 | 1390.1 | 1419.7 | 97.3 | 9749.5 |
| CD34T+ | 1550 | 341.0 | 28.2 | 50.0 | 50.0 | 480.0 |
| CD34T+ | 1800 | 813.0 | 108.0 | 489.1 | 14.8 | 2536.5 |
| CD34T+ | 1571 | 403.7 | 17.2 | 10.7 | 19.3 | 1140.2 |
| CD34T+ | 1618 | 244.8 | 2.2 | 15.2 | 17.3 | 1148.0 |
| CD34T+ | 2372 | 490.0 | 10.4 | 0.0 | 12.5 | 942.4 |
| CD34T+ | 2381 | 739.6 | 13.8 | 4.0 | 7.9 | 765.3 |

| | | No. of cells/ $10^6$ cells in spleen at w2 | | | | |
|---|---|---|---|---|---|---|
| Model | Mouse ID | hu-CD3+ | hu-CD4+ | hu-CD8+ | hu-CD19+ | mu-CD45+ |
| CD34 | 1306 | 17114.5 | 7374.7 | 8082.3 | 161409.3 | 71989.1 |
| CD34 | 1321 | 37657.2 | 31768.0 | 4971.5 | 6183.5 | 196772.1 |
| CD34 | 1300 | 37727.8 | 23376.3 | 11706.6 | 443152.9 | 51384.4 |
| CD34 | 1337 | 14563.5 | 12353.1 | 109.0 | 9113.9 | 588010.9 |
| CD34 | 1576 | 30366.1 | 23547.6 | 5864.8 | 62731.9 | 251566.1 |
| CD34 | 1802 | 16274.3 | 9329.9 | 5225.0 | 175796.5 | 399541.0 |
| CD34 | 1548 | 67005.2 | 30973.4 | 23273.6 | 1525.3 | 145455.2 |
| CD34 | 1620 | 31461.8 | 28440.5 | 555.1 | 71886.5 | 199539.6 |
| CD34 | 2371 | 3950.1 | 1743.0 | 2049.7 | 94541.7 | 64010.2 |
| CD34 | 2382 | 11367.4 | 4194.6 | 6204.8 | 187007.0 | 51005.0 |
| CD34T+ | 1344 | 484012.0 | 205888.7 | 175039.3 | 210726.7 | 10652.7 |
| CD34T+ | 1317 | 616356.2 | 289125.8 | 246492.7 | 34099.6 | 181182.1 |
| CD34T+ | 1320 | 305262.1 | 120994.2 | 145051.5 | 417.8 | 131654.6 |
| CD34T+ | 1550 | 285604.5 | 96094.8 | 157161.2 | 43804.5 | 55971.5 |
| CD34T+ | 1800 | 90674.1 | 22217.6 | 49175.3 | 9393.8 | 2396.4 |
| CD34T+ | 1571 | 169227.4 | 126501.9 | 37091.7 | 16658.4 | 224019.2 |
| CD34T+ | 1618 | 35639.5 | 15070.2 | 16173.8 | 4.2 | 380874.6 |
| CD34T+ | 2372 | 22477.2 | 14118.3 | 7700.5 | 55236.5 | 43175.4 |
| CD34T+ | 2381 | 21337.2 | 11751.2 | 5853.4 | 236240.1 | 40745.5 |
